# ImmuniT Platform for Improved Neoantigen Prediction in Lung Cancer

**DOI:** 10.1101/2025.03.17.643783

**Authors:** Stephanie J. Hachey, Alexander G. Forsythe, Hari B. Keshava, Christopher C.W. Hughes

## Abstract

1

**Introduction:** Lung cancer remains the leading cause of cancer-related deaths, with most patients presenting with advanced, treatment-resistant disease. While immunotherapy has improved outcomes for some, most patients fail to mount an effective immune response due to inadequate tumor recognition. Neoantigen-based therapies offer a promising approach to personalized immunotherapy, but current discovery methods can miss immunogenic targets, particularly those with low or heterogeneous expression. To address this, we developed the ImmuniT platform, which enhances neoantigen identification by amplifying patient-specific targets from primary tumor samples, improving prediction accuracy for more precise immunotherapy.

**Methods:** Patients with lung cancer were recruited under an IRB-approved protocol, and freshly resected tumor tissue and matched blood samples were collected. Tumors were processed into single-cell suspensions, enriched for EpCAM+ epithelial cells, and treated to enhance neoantigen expression. Peripheral blood and tumor-infiltrating lymphocytes were co-cultured with cancer cells to expand neoantigen-reactive T cells. The nextneopi pipeline integrated tumor mutational burden (TMB), HLA typing, and transcriptomic data to predict immunogenic targets. MHC:epitope complexes were validated via tetramer staining to identify patient-derived, neoantigen-specific T cells.

**Results:** The ImmuniT platform demonstrated superior neoantigen prediction and T cell activation in vitro compared to conventional methods across five NSCLC patients. In one patient, it identified two neoantigens missed by standard approaches, which were validated based on their ability to stimulate tumor-infiltrating and peripheral blood lymphocytes. Across all tested samples, the platform identified a broader spectrum of immunogenic targets. These findings highlight its potential to enhance neoantigen discovery and improve personalized immunotherapy strategies.

**Conclusion:** Our findings indicate that the ImmuniT platform improves neoantigen detection in NSCLC by identifying a wider range of tumor-specific antigens, including those overlooked by conventional methods. By expanding the pool of targetable neoantigens, this technology has the potential to enhance T cell activation and optimize immunotherapy. The ImmuniT platform represents a promising advancement towards more effective, personalized treatment strategies for lung cancer patients, particularly those who do not respond to current immunotherapies.

## 2 Introduction

Lung cancer remains the leading cause of cancer-related deaths in the US and globally, with 85% of patients presenting with advanced, treatment-resistant disease [1]. Non-small cell lung cancer (NSCLC), the predominant subtype, is characterized by a high tumor mutational burden (TMB), which has been associated with improved responses to immunotherapy [2]. However, despite the success of immune checkpoint inhibitors (ICIs), only 20–25% of patients derive significant clinical benefit, largely due to insufficient immune recognition of tumors [3, 4]. NSCLC’s heterogeneity and genetic instability drive both disease progression and therapeutic resistance, yet these same factors provide an opportunity to develop patient-specific neoantigen-targeted therapies.

Neoantigens, which arise from tumor-specific somatic mutations, play a critical role in anti-cancer immune responses by eliciting tumor-reactive T cell activity [5]. Early clinical trials have demonstrated that immunotherapy can induce neoantigen-specific immune responses in NSCLC patients [6–9]. However, a major challenge remains: while NSCLC tumors may harbor an average of 75 neoantigens, these are often heterogeneously expressed within the tumor, limiting their therapeutic potential [10, 11]. Current neoantigen discovery approaches struggle to identify highly immunogenic targets, particularly those expressed at low levels or restricted to rare tumor cell populations, such as cancer stem cells [12]. This limitation allows the cancer to evade immune attack and develop resistance to therapy.

Despite advances in sequencing, bioinformatics, and mass spectrometry, the identification of tumor-specific neoantigens remains challenging [13, 14]. Shared antigens across different cancers have shown limited efficacy, shifting focus toward patient-specific tumor antigens [4]. While neoantigen-reactive T cell therapy has demonstrated promising results, its full therapeutic potential hinges on the accurate identification of highly immunogenic targets. To address this, we have developed the ImmuniT platform, which surpasses conventional methods by detecting both abundant and low-level neoantigens. This platform amplifies patient-specific neoantigens directly from primary tumor samples, leading to improved identification and enhanced neoantigen prediction for personalized immunotherapies.

## 3 Results and Discussion

### 3.1 Patient Characteristics and Tumor Stage

Patients who consented to donate tissue and blood were enrolled in this IRB-approved study. Freshly collected lung cancer tissue, in excess of clinical need, along with matched blood samples, were processed into single-cell suspensions (Figure 1a). To enhance neoantigen detection, cancer cells were treated to amplify neoantigen expression, generating a personalized roster of neoantigen predictions for each patient. Additionally, non-primed T cells were incorporated into the platform to directly enrich for neoantigen-reactive T cells. Figure 1b presents clinical and molecular data for five NSCLC patients, demonstrating substantial variability in cancer subtype, tumor stage, and molecular characteristics. Among the five patients, four are female and one is male, with ages ranging from 49 to 84 years and racial/ethnic backgrounds including Vietnamese, White, and Hispanic individuals. Smoking history varies, with three patients identified as never-smokers, while two (patients 16 and 23) have smoking histories of 10 and 25 pack-years, respectively. Disease stage ranges from IA3 to IIIA, indicating varying tumor progression. Notably, patients 004 and 022 exhibited lymph node metastasis.

**Fig. 1.**
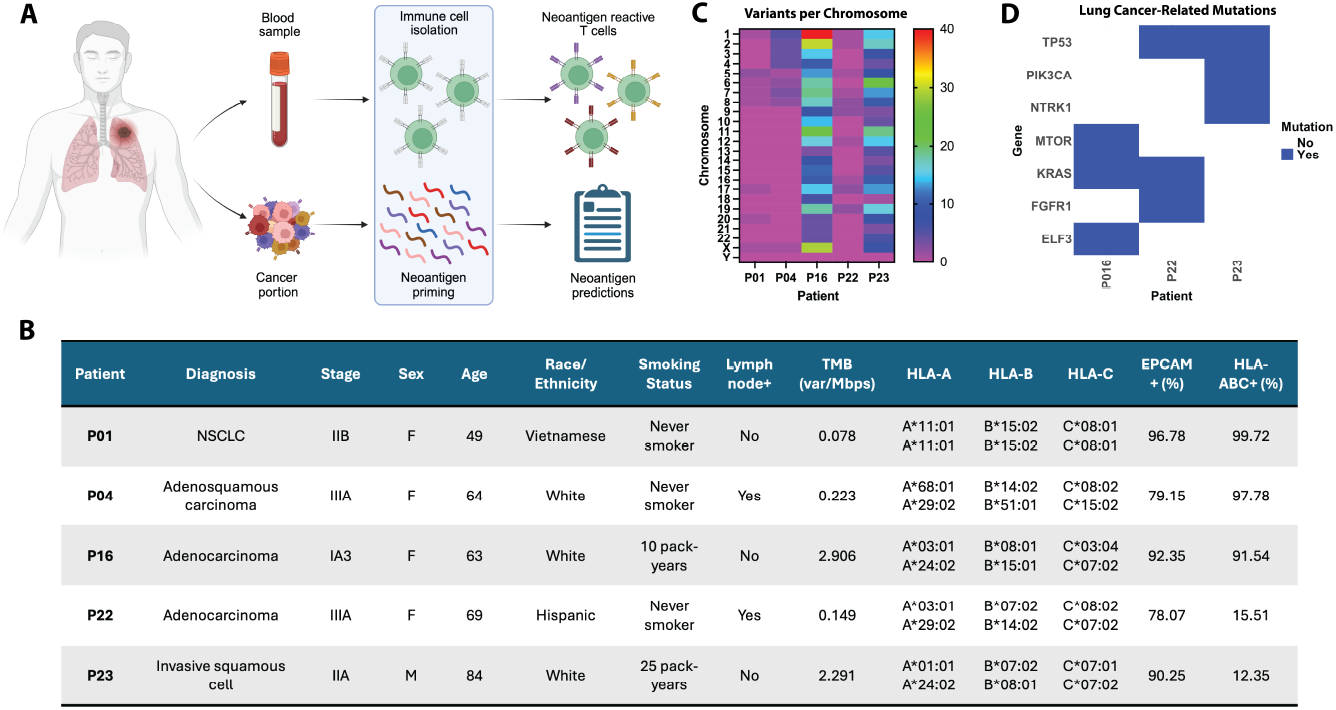
Patient and tumor characteristics. A) Schematic showing the workflow of specimen collection, live cell processing, immune cell activation and neoantigen prediction. B) Table with patient and tumor characteristics. C) Heatmap showing variants per chromosome for each patient. D) Plot showing mutation status for lung cancer-related mutations.

HLA genotyping reveals distinct allele combinations for HLA-A, HLA-B, and HLA-C across patients, which can influence antigen presentation and immune recognition. Additionally, expression levels of EpCAM and HLA-ABC highlight notable molecular differences. EpCAM positivity ranges from 78.07% to 96.78%, while HLA-ABC expression is highly variable, with patients 23 and 22 exhibiting only 12.35% and 15.51% positivity, respectively, in contrast to the high expression observed in patients 1 (99.72%), 4 (97.78%), and 16 (91.54%) (Figure 1b). HLA expression and its complexing with neoepitopes are essential for TCR-mediated recognition and tumor cell elimination [15]. Consequently, low HLA expression facilitates immune evasion and diminishes the effectiveness of immunotherapy. Tumor mutational burden (TMB), measured in variants per megabase, spans from 0.078 to 2.906 across patients. As shown in Figure 1b, patients 016 and 023, both former smokers, exhibit higher TMB compared to non-smokers, reinforcing the established link between smoking history and increased mutational load [16]. However, even among former smokers, TMB remained below the NSCLC median of 9.8 mutations/Mb [17], suggesting significant variability in mutation-driven antigenicity. The heatmap further highlights inter-patient differences in mutation distribution across chromosomes, with former smokers displaying a broader range of mutations, indicative of greater genomic instability (Figure 1c).

Figure 1d further provides insights into lung cancer-related mutations, revealing differences in oncogenic alterations between patients. P22 and P23 harbor mutations in TP53, a key tumor suppressor gene, while P16 and P22 carry mutations in KRAS, a common driver in lung cancer [18]. P16 also has mutations in MTOR and ELF3, P22 in FGFR1, and P23 in PIK3CA and NTRK1, genes involved in tumor growth and survival [19]. Interestingly, patients 1 and 4 lack mutations in these well-known oncogenic drivers, with an alternative mutational landscape contributing to tumor progression. These findings highlight the genetic heterogeneity of NSCLC, emphasizing the need for personalized therapeutic approaches based on individual mutation profiles.

### 3.2 T Cell Activation and Neoantigen Prediction with the ImmuniT Platform

To evaluate the ImmuniT platform’s ability to enhance T cell priming and activation, we analyzed patient-derived T cells from individuals with high MHC class I expression (HLA-ABC, assessed via flow cytometry). For Patient 001 (P01), pre-treatment with the ImmuniT platform significantly enhanced T cell proliferation and activation, surpassing the response observed in non-treated (standard) primed T cells (Figure 2a). In contrast, unstimulated peripheral blood lymphocytes (PBLs) exhibited minimal proliferation, while CD3/CD28 agonist (TRANSACT)-stimulated T cells showed nonspecific activation. Notably, IFN-y secretion was markedly elevated in ImmuniT-treated T cells, confirming robust immune activation and highlighting the platform’s potential to enhance immune responses.

**Fig. 2.**
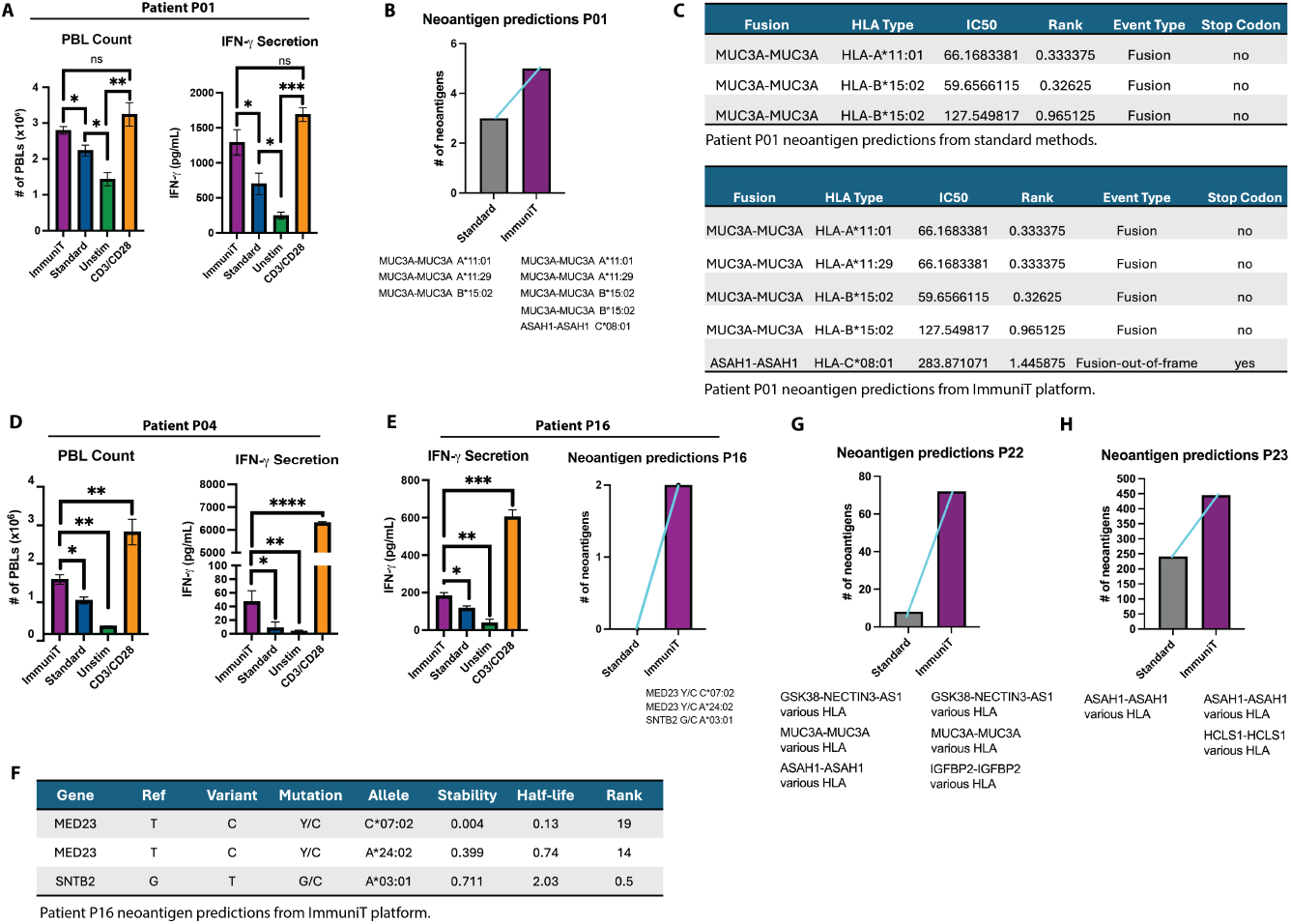
ImmuniT platform shows improved T cell activation and neoantigen detection compared to standard methods. A) Patient P01 PBL count and IFN-y secretion in each condition. B) Plot shows that standard methods detected 3 neoantigens for patient 01 lung cancer specimen, whereas ImmuniT platform detected 5 neoantigens. Neoantigens are indicated below column. C) Table showing characteristics of fusion neoantigens detected by standard methods and ImmuniT platform for patient 01 lung cancer. D) Patient P04 PBL count and IFN-y secretion in each condition. E) Patient P16 PBL IFN-y secretion in each condition. Plot shows that standard methods did not detect any neoantigens for patient 16 lung cancer specimen, whereas ImmuniT platform detected 2 neoantigens. Neoantigens are indicated below column. F) Table showing characteristics of neoantigens detected by ImmuniT platform for patient 16 lung cancer. G) Plot shows that standard methods detected 10 neoantigens for patient 22 lung cancer specimen,, whereas ImmuniT platform detected 70 neoantigens. Neoantigen classes are indicated below columns. H) Plot shows that standard methods detected 225 neoantigens for patient 23 lung cancer cells, whereas ImmuniT platform detected 450 neoantigens. Neoantigens are indicated below columns.

To determine whether the ImmuniT platform improves neoantigen detection, we integrated next-generation sequencing (NGS) and a bioinformatics pipeline to analyze primary NSCLC tumors from patients P01, P16, P22, and P23. For Patient P01, standard methods predicted three gene fusion-derived neoantigens (MUC3A-MUC3A epitopes, presented by various HLA types with differing stability and affinity)(Figure 2b,c). In contrast, the ImmuniT platform identified two additional fusion neoantigens: an alternative MUC3A fusion and an ASAH1-ASAH1 out-of-frame fusion (Figure 2b). Features of these neoantigens, including IC50 values and HLA binding rank, are detailed in Figure 2c, where higher IC50 and rank indicate stronger stability.

A similar increase in T cell activation was observed in Patient 004 (P04), where ImmuniT pre-treatment led to a significant boost in T cell proliferation compared to standard priming (Figure 2d), further demonstrating the platform’s ability to enhance immune responsiveness. Similarly, Patient 016 (P16) exhibited a notable increase in IFN-y secretion following ImmuniT treatment (Figure 2e). For P16, the ImmuniT platform also uncovered two novel neoantigens, MED23 and SNTB2, both implicated in lung cancer progression [20, 21]. These neoantigens were undetectable using conventional methodologies, underscoring ImmuniT’s ability to reveal tumor antigens with potential therapeutic relevance. Characterization confirmed HLA specificity, stability, and functional properties, supporting their potential for immunotherapy (Figure 2f). For Patient 022 (P22), ImmuniT detected fusion-derived neoantigens including GSK38-NECTIN3-AS1, MUC3A-MUC3A, and ASAH1-ASAH1, with 10 identified using standard methods versus 70 with ImmuniT. Additionally, IGFBP2-IGFBP2 emerged as a unique ImmuniT-identified target (Figure 2g). IGFBP2 (Insulin-like Growth Factor Binding Protein 2) acts as an oncogene, promoting tumor growth, migration, and invasion, and it can serve as a potential therapeutic target and biomarker [22]. In Patient 023 (P23), 225 neoantigens (ASAH1-ASAH1, various HLA types) were predicted by standard methods, whereas 450 neoantigens (ASAH1-ASAH1 and HCLS1-HCLS1, various HLA types) were identified using ImmuniT (Figure 2h, Supplemental Tables). The ASAH1 gene, which encodes the enzyme acid ceramidase, has been implicated in cancer progression [23]. Collectively, these findings suggest that the ImmuniT platform enhances antigen-specific detection and T cell priming, a critical factor for effective cancer immunotherapy.

### 3.3 Functional Validation of Novel Neoantigens

To validate the functional avidity of T cells from patient 16, who had a high mutational burden and high expression of HLA-ABC, against the predicted neoantigens, we employed a cloning and expression system to investigate T cell receptor (TCR) specificity in TILs from patient P016 (Figure 3a). Custom-designed oligonucleotides targeting MED23 and SNTB2 were integrated into a lentiviral plasmid and used to generate lentiviruses, enabling the stable transduction of autologous fibroblasts to overexpress the neoantigens (Figure 3b,c). Transduction efficiency was confirmed via fluorescence markers, ensuring robust antigen presentation. Co-culture experiments with neoantigen-expressing fibroblasts and patient-derived TILs revealed a 3.84% frequency of MED23-specific T cells, as confirmed by flow cytometry and peptide-MHC (pMHC) tetramer staining (Figure 3d). No SNTB2-specific TILs were detected (Figure 3e). However, analysis of peripheral blood lymphocytes (PBLs) demonstrated the presence of T cells reactive to both neoantigens (Figures 3f,g), suggesting that neoantigen-specific T cell populations may differ between peripheral blood and the tumor microenvironment. This observation could have significant implications for neoantigen-targeted immunotherapy, as circulating T cells might serve as a reservoir for antigen-specific immune responses.

**Fig. 3.**
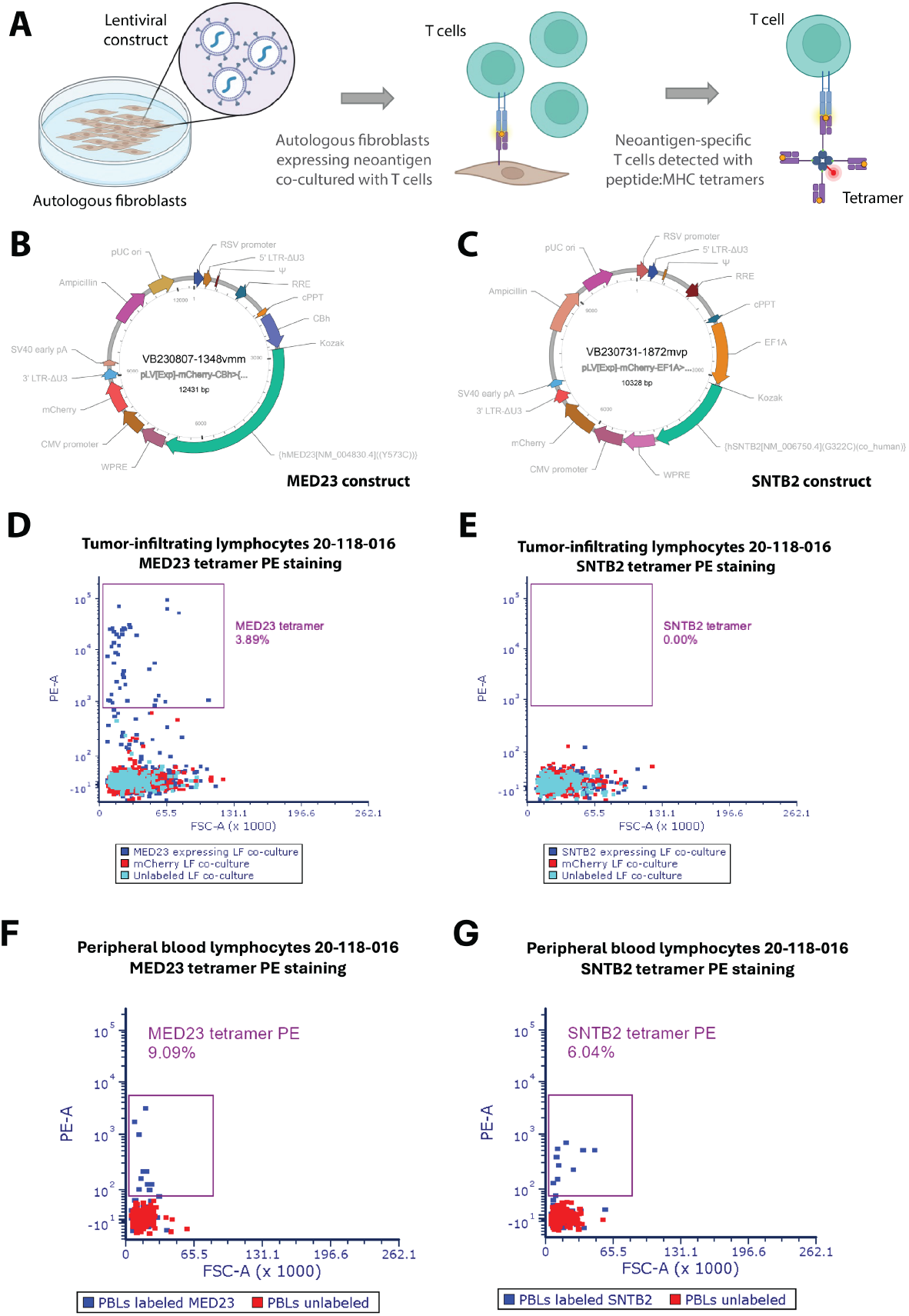
The ImmuniT platform enhances the identification of neoantigens for patient 016, as shown by the expansion of neoantigen-specific tumor-infiltrating lymphocytes detected through peptide:MHC tetramers. (A) Schematic showing workflow for assessing TCR specificity. Autologous fibroblasts are transduced with a lentiviral construct containing the neoantigen of interest, then co-cultured with patient-specific T cells. T cells with TCR specific to neoepitope:MHC presented on the cell surface are enriched and detected via peptide:MHC tetramer staining and flow cytometry. (B) Plasmid containing neoantigen construct and fluorescent reporter for MED23 and (C) SNTB2, putative neoantigens detected by ImmuniT platform for patient 016. (D) Flow cytometric analyses of tumor-infiltrating lymphocytes (TILs) from patient 016 co-cultured with MED23 expressing autologous lung fibroblasts (LF) in addition to TILs co-cultured with control mCherry expressing LF and unlabeled LF for MED23 and (E) SNTB2 peptide:MHC tetramer staining. Note that 3.89% of TILs co-cultured with MED23 expressing LF show TCR specificity for MED23, whereas 0% of TILs co-cultured with SNTB2 expressing LF show TCR specificity for SNTB2. (F) Flow cytometric analyses of peripheral blood lymphocytes (PBLs) from patient 016 co-cultured with MED23 expressing autologous lung fibroblasts (LF) in addition to PBLs unlabeled LF for MED23 and (G) SNTB2 peptide:MHC tetramer staining. Note that 9.09% of PBLs show TCR specificity for MED23, whereas 6.04% of PBLs show TCR specificity for SNTB2.

### 3.4 Implications for Personalized Immunotherapy in NSCLC

Our study reveals that conventional neoantigen identification methods may miss functionally relevant targets, potentially limiting the efficacy of personalized cancer vaccines and adoptive T cell therapies. By utilizing the ImmuniT platform, we achieved enhanced T cell activation and proliferation, improved neoantigen detection, and functional validation of neoantigen-specific T cells, uncovering differential distribution of neoantigen-specific T cells between peripheral blood circulating lymphocyte populations and tumor resident compartments. In our analysis of patient P016, flow cytometry and pMHC tetramer assays demonstrated that while tumor-infiltrating lymphocytes (TILs) contained MED23-specific T cells, no SNTB2-specific T cells were detectable within the tumor microenvironment. In contrast, both MED23- and SNTB2-reactive T cells were present in peripheral blood lymphocytes (PBLs). This compartmental bias suggests that peripheral blood may serve as a more accessible source of neoantigen-specific T cells, but also raises concerns about potential biases introduced during T cell isolation, culture, and expansion.

These findings have significant implications for both vaccine-based and cell-based immunotherapies. For cell-based therapies, selecting the optimal starting population is crucial for approaches such as TIL therapy or adoptive transfer of neoantigen-specific T cells [8, 11, 24, 25]. The underrepresentation or functional suppression of certain tumor-reactive clones within the tumor microenvironment may limit therapeutic efficacy, and current expansion protocols might preferentially amplify T cells with superior growth characteristics rather than those with the highest tumor reactivity [26, 27]. For vaccine-based therapies, therapeutic vaccines depend on the priming and expansion of endogenous neoantigen-reactive T cells to generate a robust anti-tumor response [9, 28]. By enhancing neoantigen identification, the ImmuniT platform could improve vaccine efficacy by expanding rare but functionally significant T cell clones before antigen exposure. Given the observed biases in neoantigen-specific T cell distribution, understanding the compartmental distribution of neoantigen-specific T cells is therefore essential to optimizing immunotherapy and improving patient outcomes. Additionally, the selective pressures within the tumor microenvironment may deplete highly functional T cell clones, further complicating efforts to harness an effective anti-tumor response.

By combining next-generation sequencing with functional validation assays, our study demonstrates that the ImmuniT platform can identify rare but immunologically relevant neoantigens, even in NSCLC tumors with a low tumor mutational burden. A key strength of the platform is its HLA-agnostic approach, which enables the identification and validation of neoantigens without being constrained by HLA restrictions. Moving forward, efforts should prioritize refining neoantigen-specific T cell expansion protocols and optimizing therapy designs to address compartmental biases and functional constraints. By enhancing T cell activation, improving neoantigen detection, and circumventing the challenges associated with low TMB, the ImmuniT platform establishes a powerful foundation for developing highly personalized and broadly accessible immunotherapies.

## 4 Methods

### 4.1 Tissue and Blood Collection

Surgically resected NSCLC tissue and matched blood samples were collected from 20 patients under an IRB-approved protocol at the University of California, Irvine. All experimental procedures were conducted in accordance with institutional guidelines and regulations, with informed consent obtained from all participants or their legal guardians. Adjacent normal lung tissue, resected beyond tumor margins to ensure complete disease removal, was also included in the study. Blood samples were collected in 7.5 mL CPT Mononuclear Cell Preparation tubes containing sodium heparin and Ficoll. Surgical resections were collected in a conical tube containing transport media and transported on ice within 2 hours of collection. The transport media consists of DMEM (4.5 g/L glucose, sodium pyruvate, and L-glutamine; Cat# 10-013-CV), serum-free with 1x P/S, antimitotic/antimycotic (Anti-anti 100x, A5955, Millipore), and 10 *µ*M ROCK inhibitor (Y-27632, StemCell Technologies, Cat# 72302).

### 4.2 Tissue Processing and Dissociation

NSCLC and normal lung tissues were subjected to mechanical dissociation using scalpels to finely mince the samples. As the pieces become smaller, gentle pipetting with a P1000 facilitates the liberation of cell clusters suitable for direct plating. These liberated cells should be collected, centrifuged, and plated. These liberated cells were centrifuged at 300 × g for 10 minutes, the supernatant removed, and the pellet washed twice with HBSS before plating into an ultra-low attachment dish containing LCIC-supplemented media. To remove immune cells, an additional centrifugation step at 200 × g for 3 minutes in 50 mL HBSS was performed, with immune cells remaining in the supernatant. If dissociation was incomplete, the second step was initiated. Remaining chunks of minced tissue were then incubated in 1x triple enzyme digestion mix (prepared from 10x stock containing 1 g Collagenase, 20,000 units DNase, and 100 mg Hyaluronidase in 100 mL HBSS) supplemented with 10 *µ*M ROCK inhibitor. Samples were shaken at room temperature in 30-minute intervals, with periodic assessment for adequate digestion. Liberated single cells were either plated for culture, frozen for later use, or subjected to further processing as needed. CD3 microbeads were used to isolate tumor-infiltrating lymphocytes (TILs), which were then frozen. CD45 microbeads were used to remove other immune cells, which were also frozen following the microbead protocol. Primary lung fibroblasts were isolated by plating cells onto tissue-culture treated flasks in DMEM 10% FBS and confirmed by staining with PDGFRa and PDGFRb. Cell viability was assessed using flow cytometry with PI. To confirm the tumor origin of the sample, FACS analysis was performed using antibodies targeting EpCAM.

### 4.3 Cell culture and co-culture assays

We first isolated cancer cells by sorting dissociated tissue for EpCAM+ cells via flow cytometry and expanding NSCLC cell numbers for no more than a week in specialized cancer-initiating cell medium containing. Normal lung epithelial cells were grown in lung cancer medium supplemented with Wnt3a and Noggin. MHC class I expression was also assessed by flow cytometry (HLA-ABC antibody). From 5 of the patient-derived NSCLC specimens, several methods were tested to improve expression and accumulation of neoantigens, remove tolerogenic molecules, and enhance vaccine immunogenicity. Autologous peripheral blood lymphocytes were isolated and subjected to *ex vivo* expansion in TexMACS medium supplemented with IL-2. We looked to see if we had generated activated T cells through our procedure by performing lymphocyte proliferation assays and interferon-gamma ELISAs.

### 4.4 Neoantigen prediction pipeline

To identify neoantigen candidates, we utilized the nextNEOpi pipeline to process raw DNA and RNA sequencing data from paired tumor-normal samples, including those subjected to ImmuniT priming [14]. nextNEOpi is a Nextflow-based computational framework that integrates multiple stages, including data preprocessing, variant identification, HLA typing, neoantigen prediction, and quantification of tumor immunogenicity-related features. For class-I neoepitope prediction, nextNEOpi employs pVACseq, which incorporates netMHCpan, netMHCIIpan, and mhcflurry, all of which leverage machine-learning algorithms to predict MHC-presented neoantigens. Fusion neoepitopes were identified using NeoFuse. Raw whole-exome sequencing (WES) and RNA sequencing data underwent quality control to remove low-quality reads, adapter sequences, and contaminants. Somatic variants were identified by comparing tumor and normal WES data, and HLA typing was performed to determine the MHC molecules expressed by tumor cells. Neoantigen candidates were then predicted based on somatic variant-derived peptides capable of MHC class I presentation on the tumor cell surface. Predictions were confirmed against the TESLA data results outlined in [13].

### 4.5 Neoantigen Expression Assays

Lentiviral vectors encoding neoantigens were designed and constructed by Vector-Builder. To generate lentiviruses, HEK293T cells were transfected with the lentivector and packaging plasmids using Lipofectamine 2000, following the manufacturer’s protocol (Thermo Fisher Scientific). Viral supernatants were collected at 24 and 48 hours post-transfection and concentrated by incubating with a 50% PEG solution in sterile PBS for 24 hours. The resulting viral precipitate was pelleted by centrifugation at 2,500 rpm for 20 minutes, after which the supernatant was discarded, and the viral pellet was resuspended in PBS. Viral titers were determined using a P24 ELISA kit. For transduction, primary patient-derived lung fibroblasts at 50% confluence were exposed to the concentrated lentivirus at a multiplicity of infection (MOI) of 1 in the presence of 8 *µg/mL* polybrene.

### 4.6 Tetramer staining and flow cytometry

Custom MHC-peptide tetramers were designed and produced by BioLegend to detect antigen-specific T cells in non-small cell lung cancer (NSCLC) patient samples. The tetramers used in this study included HLA-A*24:02, MED23 (TYSRLLVCM-PE) and 10 *×* HLA-A*03:01, SNTB2 (ATSTAGCSK-PE). Peripheral blood mononuclear cells (PBMCs) or tumor-infiltrating lymphocytes (TILs) were first enriched for T cells using CD3 MicroBeads (Miltenyi Biotec) and magnetic-activated cell sorting (MACS) to isolate CD3^+^ T cells. The purified cells were resuspended in staining buffer (PBS + 2% FBS) at 1 × 10^6^ cells per 100 *µ*L and incubated with PE-labeled tetramers at room temperature for 30 minutes in the dark. Flow cytometry analysis was performed on a BD Fortessa high-parameter flow cytometer, and data were analyzed using FCS Express software. Tetramer-positive T cells were identified by gating on live, single cells, with proper fluorescence compensation and gating strategies applied using fluorescence-minus-one (FMO) controls and unstained samples.

### 4.7 Statistical Analysis

Each experimental group consisted of three independent biological replicates, with each experiment performed in technical triplicates. For comparisons across multiple groups, one-way ANOVA was conducted to assess overall variance and estimate experimental error. Pairwise comparisons between treatment and control groups were performed using a two-tailed t-test, with statistical significance set at *p <* 0.05.

## Supplementary information

Supplemental Tables are included.

## Acknowledgements

Biorender was used to generate Figure 1a and Figure 3a. This work was supported by the National Institutes of Health, National Cancer Institute, Department of Defense, and National Center for Advancing Translational Sciences through the following grants: UG3/UH3 TR002137, R61/R33 HL154307, 1R01CA244571, 1R01 HL149748, U54 CA217378(CCWH) and 75N91022C00004, W81XWH2110393, and TL1 TR001415 (SJH).

## Declarations

All patients enrolled in this IRB-approved study provided informed consent, and all research procedures adhere to ethical guidelines and regulatory standards. SJH and AGF have an equity interest in ImmunoTarget Therapeutics, Inc., which is commercializing some of the technology described in this paper. The terms of this arrangement have been reviewed and approved by the University of California, Irvine in accordance with its conflict of interest policies.

